# Molecular Basis for N-terminal Alpha-Synuclein Acetylation by Human NatB

**DOI:** 10.1101/2020.05.11.089318

**Authors:** Sunbin Deng, Buyan Pan, Leah Gottlieb, E. James Petersson, Ronen Marmorstein

**Author notes:** Correspondence (R.M.).

## Abstract

NatB is one of three major N-terminal acetyltransferase (NAT) complexes (NatA-NatC), which co-translationally acetylate the N-termini of eukaryotic proteins. Its substrates account for about 21% of the human proteome, including well known proteins such as actin, tropomyosin, CDK2, and α-synuclein (αSyn). Human NatB (hNatB) mediated N-terminal acetylation of αSyn has been demonstrated to play key roles in Parkinson’s disease pathogenesis and as a potential therapeutic target for hepatocellular carcinoma. Here we report the cryo-EM structure of hNatB bound to a CoA-αSyn conjugate, together with structure-guided analysis of mutational effects on catalysis. This analysis reveals functionally important differences with human NatA and *Candida albicans* NatB, resolves key hNatB protein determinants for αSyn N-terminal acetylation, and identifies important residues for substrate-specific recognition and acetylation by NatB enzymes. These studies have implications for developing small molecule NatB probes and for understanding the mode of substrate selection by NAT enzymes.

## Introduction

N-terminal acetylation (NTA) is an irreversible protein modification that predominantly occurs co-translationally on eukaryotic nascent chains emerging from the ribosome exit tunnel [1]. This modification occurs on ∼ 80% of human proteins [2] and impacts several protein functions including complex formation [3–7], protein localization [8–12], and the N-end rule for protein degradation [13–15]. NTA is catalyzed by a group of enzymes called N-terminal acetyltransferases (NATs) which belong to the Gcn5-related N-acetyltransferase (GNAT) family of enzymes. In humans, seven NATs have been identified to date: NatA-F, and NatH [1]. NatA, NatB, NatC are responsible for the majority of NTA [16]. NatA and NatB are heterodimeric complexes, each with a distinctive catalytic subunit (NAA10 in NatA, and NAA20 in NatB) [17] and a unique auxiliary subunit (NAA15 in NatA, and NAA25 in NatB) [18]. NatC is a heterotrimeric complex that contains a catalytic subunit (NAA30) and two auxiliary subunits (NAA35 and NAA38) [19]. Structural and biochemical studies of NatA reveal that the auxiliary subunit plays an important allosteric role in substrate binding specificity and catalysis[20] and also contributes to ribosome binding during translation [21, 22]. NatA, NatB and NatC exhibit distinguishable substrate specificity mainly toward the first two residues of the nascent chain N-termini. NatA accounts for 38% of the human N-terminal acetylome, acetylating small N-terminal residues after removal of the initial methionine residue [23], while NatB modifies N-termini with sequences of MD-, ME-, MN- and MQ- [24, 25]. NatC/E/F have overlapping substrates, acting on N-terminal methionine when it is followed by several residues excluding D, E, N and Q [23, 26–31]. When another catalytic subunit (NAA50) binds to NatA, a dual enzyme complex NatE is formed [32, 33], with catalytic crosstalk between NAA10 and NAA50 [33, 34]. We recently demonstrated that both NAA50, and a protein with intrinsic NatA inhibitory activity - Huntingtin-interacting protein K (HYPK) [35–37], can bind to NatA simultaneously to form a larger tetrameric complex [33]. NatD and NatH are highly selective enzymes with restricted substrates, displaying activity toward H4/H2A and N-terminally processed actin, respectively [38–40]. The unique localization of NatF to the Golgi membrane demonstrates NTA can occur post-translationally in some cases [41].

NatB is conserved from yeast to human in both complex composition and in its substrate specificity profile [16]. In *Saccharomyces cerevisiae*, the deletion of NatB subunits produces more severe phenotypes, compared to the knockout of NatA or NatC subunits. Deletion of either NAA20 or NAA25 leads to similar phenotypes including slower growth rate, diminished mating, defects in actin cable formation, and aberrant mitochondrial and vacuolar inheritance [42]. These observations suggest that the proper function of actin and tropomyosin requires NTA by the intact NatB complex [42]. In humans, disruption of NatB (hNatB) by knockout leads to defects in proper actin cytoskeleton structure, cell cycle progression and cell proliferation [17, 43–45]. In addition, hNatB is upregulated in human hepatocellular carcinoma [43], where it has been suggested as a potential therapeutic target as silencing of this complex can block cell proliferation and tumor formation [45]. hNatB-mediated NTA of α-synuclein (αSyn) has been shown to increase αSyn stability and lipid binding, and to reduce aggregation capacity [10, 46–53]. Since αSyn is a key protein in Parkinson’s disease (PD) [54, 55], hNatB might play an indirect role in PD pathogenesis *in vivo* as supported by a recent study [56].

Compared to the comprehensive structural and biochemical characterization of NatA [20–22, 34–36], the study of NatB has been limited, particularly in humans. Recently, the crystal structure of *Candida albicans (Ca)* NatB bound to a bisubstrate CoA-peptide conjugate was determined, providing important insights into substrate specificity and NTA by caNatB [57]. However, hNAA20 and hNAA25 share only ∼40% and ∼20% sequence identity with the *Candida albicans* homologue [57], respectively. Moreover, nearly all biological studies of NatB have been conducted in *Saccharomyces cerevisiae* [58–60], Arabidopsis [61, 62], mouse [63] and human [17, 43–45] as model organisms. As a result, the mode of human NatB-mediated catalysis and αSyn-specific NatB recognition remains unresolved. In this study, we report the 3.5 Å resolution cryo-electron microscopy (cryo-EM) structure of hNatB bound to a bisubstrate CoA-αSyn conjugate, together with a structure-guided analysis of mutational effects on catalytic activity. This analysis reveals functionally important structural differences between hNaB and related NAT enzymes, as well as insights into the molecular mechanisms that define αSyn and related substrates that are recognized for hNatB-mediated N-terminal acetylation.

## Results

### hNatB is potently inhibited by a CoA-αSyn conjugate

While attempts to express recombinant hNatB in *E. coli* were unsuccessful, we found that overexpression of hNatB complex with full-length hNAA25 (residues 1-972) and C-terminally truncated hNAA20 (residue 1-163 out of 178 total residues) in baculovirus-infected Sf9 insect cells produced soluble protein that could be purified to homogeneity (**Fig. 1A**). To evaluate the activity of the recombinant hNatB, we tested it against different peptide substrates. αSyn with an N-terminal sequence of “MDVF” has been widely considered as an *in vivo* hNatB substrate [24, 64–66]. We therefore incorporated this sequence into a peptide substrate named “MDVF” for an *in vitro* acetyltransferase assay (see Methods for the full sequence). In agreement with *in vivo* studies [24, 64–66], we observed that the purified recombinant hNatB was active against this “MDVF” peptide, while no activity could be observed in the absence of either the enzyme or peptide (**Fig. 1B**). hNatB also showed no observable activity if either the first residue “M” or the first two residues “MD” in this αSyn peptide substrate was removed (**Fig. 1B**), suggesting that peptide substrate recognition by NatB is highly dependent on the first two N-terminal residues. To further confirm the substrate specificity of hNatB, we tested it against several previously identified peptide substrates for other NATs (**Fig. 1B**, see Methods for full peptide sequences): “SASE” peptide (NatA); “MLRF” peptide (NatC); “SGRG” (NatD); “MLGP” peptide (NatE). Consistent with previous results [24], hNatB is only active toward its unique canonical substrate type, displaying no overlapping activity towards other NAT substrates.

**Fig. 1.**
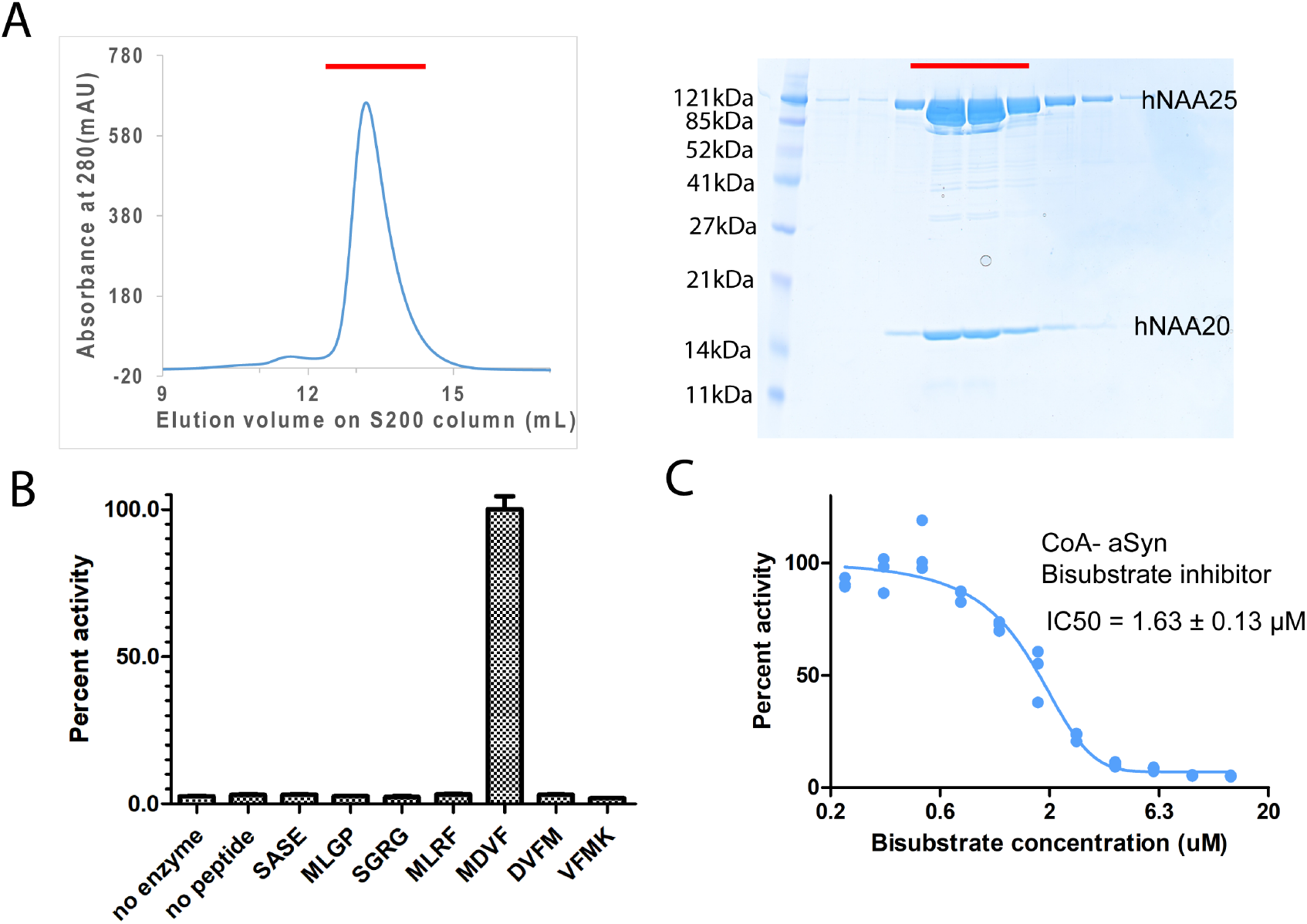
hNatB is active toward an α-Synuclein peptide and can be inhibited by a CoA-αSyn conjugate. **A**. Gel filtration elution profile of hNatB, using a Superdex S200 column. Coomassie-stained SDS-PAGE of peak fractions is reproduced to the right of the chromatograms. **B**. Comparison of hNatB activity toward different peptide substrates. All the activities are normalized to the activity of hNatB toward αSyn peptide (MDVF). **C**. Dose-response curve corresponding to the titration of CoA-αSyn conjugate (CoA-MDVFMKGLSK) into hNatB acetyltransferase reactions. The calculated IC_50_ value is indicated. Reactions were performed in triplicate; replicates are shown in the graph as vertical dots.

In order to understand the mechanism of hNatB substrate recognition, we synthesized a bi-substrate inhibitor in which the first 10 residues of αSyn are covalently linked to CoA [20] for enzymatic and structural studies. Half-maximum inhibitory concentration (IC_50_) determinations revealed that this CoA-αSyn conjugate had an IC_50_ of about 1.63 ± 0.13 μM (**Fig. 1C**), significantly lower than the K_m_ values we had determined for hNatB toward a “MDVF” peptide (45.08 ± 3.15 μM) and acetyl-CoA (47.28 ± 5.70 μM) **(Table 1)**.

**Table 1.**
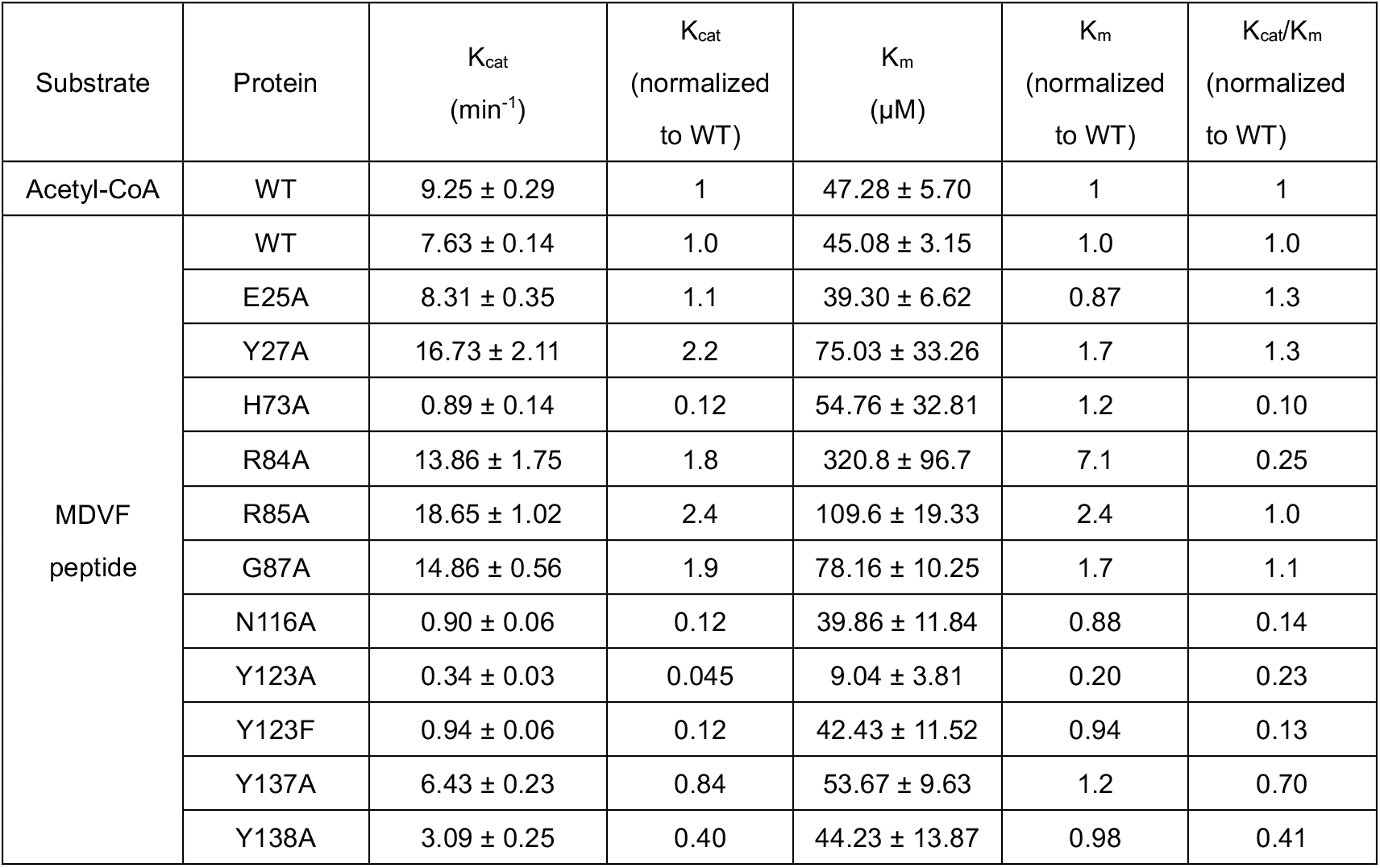
Catalytic parameter of wild-type hNatB and mutants.

### hNatB reveals potentially biologically significant structural differences with hNatA and caNatB

We performed single particle cryo-EM of hNatB in the presence of the CoA-αSyn conjugate. A 3.46 Å-resolution cryo-EM threedimensional (3D) map was determined from 982,420 particles, selected from 5,281 raw electron micrographs **(Table 2, and Fig.S1 and S2)**. The central core region of the EM map contains excellent side chain density with a local resolution of ∼ 2.5 Å, particularly around the catalytic subunit, hNAA20.

**Table 2.** Cryo-EM data collection, refinement and validation statistics.

|  |  |
| --- | --- |
| | hNatB/CoA-αSyn complex<br>EMD-21307<br>PDB: 6VP9 |
| **Data collection and processing** |  |
| Magnification | 105,000 |
| Voltage (keV) | 300 |
| Electron exposure (e/Å<sup>2</sup>) | 40 |
| Defocus range (μm) | -1.5 to -2.5 |
| Pixel size (Å) | 0.83 |
| Symmetry imposed | C1 |
| Initial particles(no.) | 1,927,675 |
| Final particles (no.) | 982,420 |
| Map resolution (Å) | 3.46 |
| FSC threshold | 0.143 |
| Map resolution range (Å) | 2.5-4.5 |
| **Refinement** |  |
| Initial model used (PDB code) | - |
| Model resolution (Å) | 3.5 |
| FSC threshold | 0.5 |
| Model resolution range (Å) | - |
| Map sharpening *B* factor (Å<sup>2</sup>) | -191.177 |
| **Model composition** |  |
| Non-hydrogen atoms | 8611 |
| Protein residues | 1077 |
| Ligands | 2 |
| ***B* factors (Å<sup>2</sup>)** |  |
| Protein | 2.51/84.68/33.39 |
| Ligand | 17.07/19.91/19.74 |
| **R.M.S. deviations** |  |
| Bonds lengths (Å) | 0.011 |
| Bond angles (°) | 1.105 |
| **Validation** |  |
| MolProbity score | 1.73 |
| Clash score | 3.98 |
| Poor rotamers (%) | 0.55 |
| **Ramachandran plot** |  |
| Favored (%) | 90.10 |
| Allowed (%) | 9.71 |
| Disallowed (%) | 0.19 |

Consistent with previous NAT structural studies, the atomic model of hNatB features a catalytic subunit, hNAA20, that adopts a canonical Gcn5-related N-Acetyltransferase (GNAT) fold [67]. Additionally, the model reveals that the auxiliary subunit hNAA25 is composed of a total of 39 α-helices among where the predicted first and second α-helices were built as poly-alanine due to a lack of resolvable side-chain density **(Fig.S3)**. The 39 α-helices can be roughly divided into three groups: an N-terminal region: α1-α8; a core region: α9-α29; and a C-terminal region: α30-α39 **(Fig. 2A and Movie 1)**. The eight helices of the N terminal region form four helical bundle tetratricopeptide repeat (TPR) motifs, which often participate in protein-protein interactions. While there are no visible contacts between the N-terminal TPR motifs and hNAA20, it is possible that this region participates in ribosome association, similar to the N-terminal region of the NAA15 auxiliary subunits of *Schizosaccharomyces pombe* [21] and *Saccharomyces cerevisiae* [22] NatA. The 21 helices of the core region also form a number of TPR motifs, which come together to form a ring that completely wraps around and extensively contacts hNAA20 within its hollow center **(Fig. 2A)**. Indeed. the interaction between hNAA20 and the TPR motifs of this core region buries a total interface area is about 2300 Å^2^. In the core region, it is noteworthy that there is a long α-helix (α28, ranging 30 residues) that traverses almost from one side of the complex to the other. The α28 helix closes the core ring structure, locking hNAA20 in position, and bridging the N- and C-terminal and regions. This is similar to the role played by α29-α30 of the hNAA15 auxiliary subunit of hNatA **(Fig. 2A and 2B)**. The C terminal region features helices that bundle together to protrude out of the plane of the core ring structure at an angle of ∼ 45° (**Fig. 2B**).

**Fig. 2.**
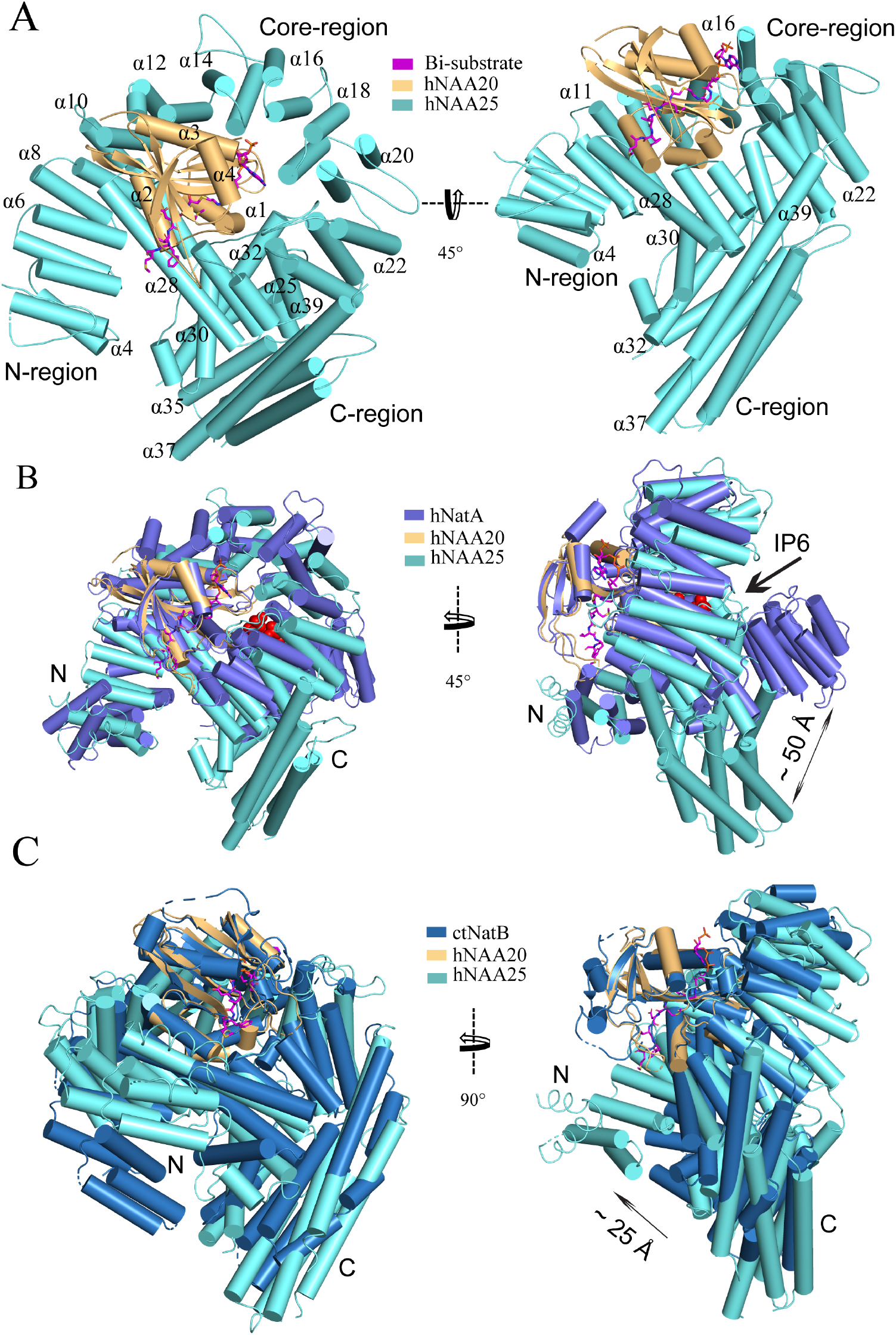
hNatB shows structural differences with hNatA and *C. albicans* NatB. **A**. hNaa20 (light orange) and hNaa25 (cyan) are shown in cartoon. The CoA-αSyn conjugate inhibitor is shown in sticks and colored as magenta. The N- and C-terminal regions are indicated as “N” and “C”, respectively. Some helices are as labeled. **B**. hNaa20 (light orange) and hNaa25 (cyan) are shown overlapped with hNatA (marine blue, PDB: 6C9M). Small molecule IP_6_ bound to hNatA is shown as surface representation (red). **C**. hNaa20 (light orange) and hNaa25 (cyan) are shown superimposed on *Ca*NatB (slate blue, PDB:5K04).

The sequence identity of the catalytic and auxiliary subunits of hNatA and hNatB are 20%, and 15 %, respectively. To understand how this translates to key structural differences, we superimposed the crystal structure of hNatA (PDB: 6C9M) with our model. Between the catalytic subunits, there is a high degree of superposition (1.151 Å root-mean-square deviation [RMSD] over 105 common C_α_ atoms), except for an additional helix, α5, on the C-terminus of hNatA-NAA10, which is absent in hNatB-NAA20 **(Fig. 2B)**. Between the auxiliary subunits, the core and N-terminal regions of both hNatA-NAA15 and hNatB-NAA25 display similar topology, although a higher degree of deviation than the catalytic subunits. The core regions both wrap around their respective catalytic subunits (8.369 Å RMSD over 262 common C_α_ atoms), while the N-terminal regions jut off to the side (7.360 Å RMSD over 55 common C_α_ atoms). In contrast, the C-terminal regions of hNatA-NAA15 and hNatB-NAA25 diverge significantly from one another. For hNA25, the C-terminal region of hNAA25 is oriented towards its N-terminal region, while the C-terminal region of hNAA15 is positioned ∼ 50Å away from the relative position of the superimposed C-terminal domain of hNAA25 **(Fig. 2B)**. The positioning of hNAA25 may serve to promote hNAA25 intra-termini communication, which is similar to the interaction of hNatA and its regulatory protein HYPK [35]. HYPK, which does not interact with hNatB, interacts with both the N- and C-terminal domains of hNatA-NAA15, potentially serving as bridge to enable closer communication between these two domains **(Fig. S4*A*)**. Recent reports have described the role of the small molecule IP_6_ (inositol hexakisphosphate) in hNatA activity, where it is found to act as “glue” between the C-terminal and core domains in hNAA15 and hNAA10 via a series of hydrogen bonds and electrostatic interactions [35, 68]. While no corresponding small molecules have been identified to play a similar role in hNatB, our model shows that this interaction is replaced by an extended loop that connects the a31 helix with the a32 helix of hNatB-NAA25. This loop, which is not present in hNatA-NAA15, appears to mediate hydrophobic interactions between hNatB-NAA25 and -NAA20, likely to serve a similar role as IP_6_ **(Fig. S4*B*)**.

We also compared the structures from human and the previously described *C. albicans* NatB (*Ca*NatB, PDB: 5K18). Although the two superimposed structures revealed a high degree of structural conservation (NAA20: 0.698 Å RMSD over 125 common C_α_ atoms; NAA25 Core region: 3.267 Å RMSD over 266 common C_α_ atoms), the N-terminal region of hNatB-NAA25 appears to overlay more closely to hNatA-NAA15 than to *Ca*NatB-NAA25 **(Fig. 2B and C)**. Compared to *Ca*NAA25, the N-terminal regions of hNatB-NAA25 and hNatA-NAA15 are positioned more closely to the peptide substrate binding sites of the respective catalytic subunits. Based on the role that the N-terminal yeast NatA-Naa15p regions play in ribosome docking [21, 22], we propose that the relative shift in position of the N-terminal regions of the human NAT auxiliary subunits, hNAA15 and hNAA25, may reflect a difference in mechanism for ribosome association and co-translational NTA in *C. albicans* compared with humans. In addition, the overlay of C terminal regions of hNAA25 and CaNAA25 displays a RMSD of 15.960 Å over 133 common C_α_ atoms. We observe that the main difference that contributes to this deviation in this region is the length of helices.

### hNAA25 and hNAA20 make intimate interactions within hNatB

The hNatB/CoA-αSyn structure reveals an extensive interaction interface between the core region of the auxiliary hNAA25 and catalytic hNAA20 subunits (**Fig. 3A**). The most intimate contact between the two proteins are made by the α28 - α29 segment of hNAA25 and almost the entire length of hNAA20 α2, creating a large hydrophobic interface (**Fig. 3B**). Residues that contribute to interaction include Thr26, Gly28, Ile29, Pro30, Leu33, Gln34, Leu36, Ala37, His38, Glu41 of hNAA20 and Lys535, His536, Ile537, Phe569, Asp576, Thr577, Tyr580, Ala584, Tyr 587, Lys592, Phe596, Phe599, Leu603 of hNAA25 (**Fig. 3B**). Another region of interaction, involving predominantly van der Waals interaction occurs between the hNAA20 β4-α3 loop and the hNAA25 α18-α19 loop. Here, residues Lys362, Pro363, Lys362-Thr367, Cys364, and Pro363 from hNAA25 and residues Arg85, Phe83, Glu82 and Arg84 from hNAA20 contribute to this small batch of hydrophobic interface (**Fig. 3C**).

**Fig. 3.**
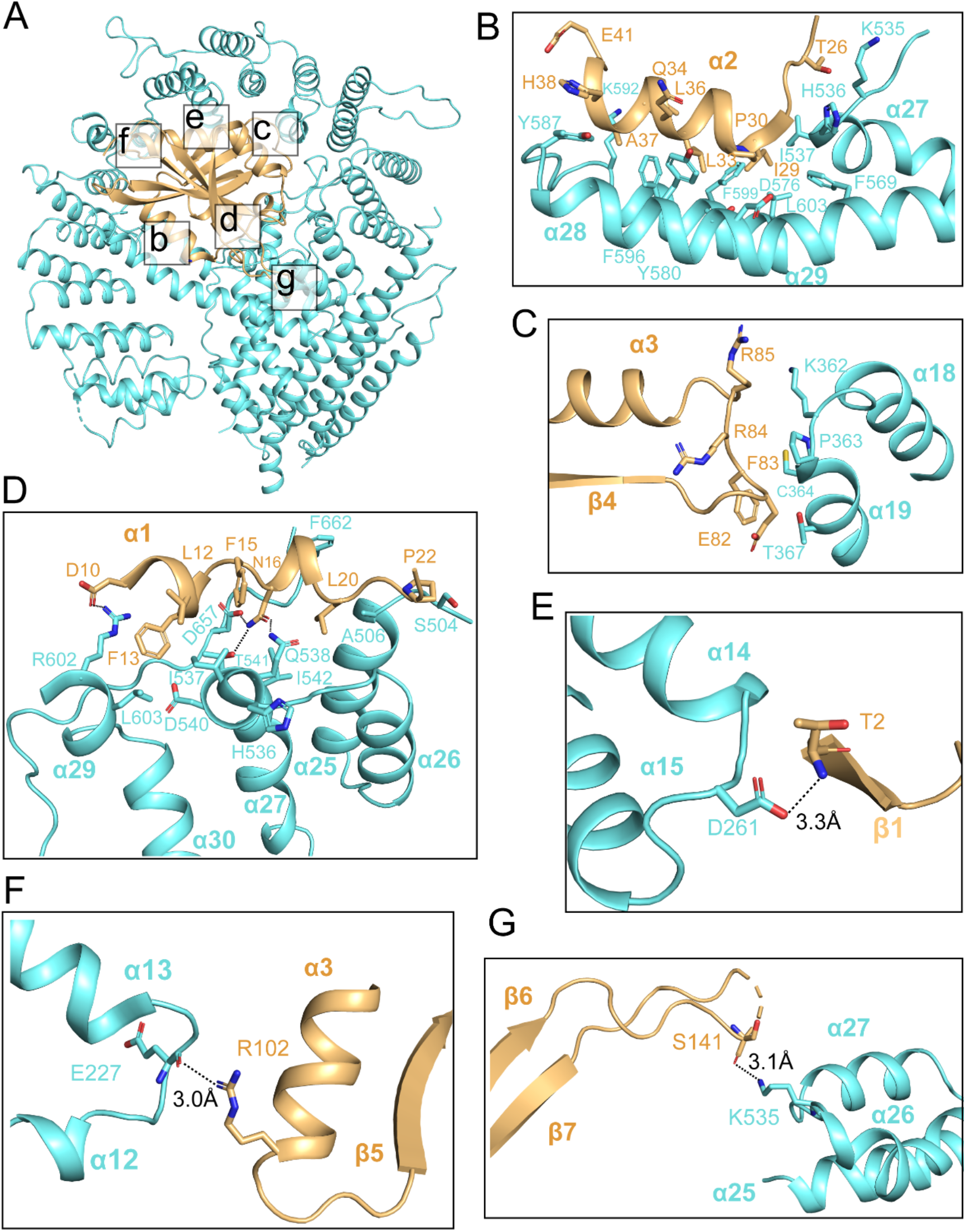
hNAA20 and hNAA25 make intimate interactions within hNatB. **A**. hNAA20 (light orange) and hNAA25 (cyan) are shown in cartoon with major associated interface denoted. **B-G**. Zoom-in views of the hydrophobic interface regions as indicated in **A**.

Additional intimate contacts between hNAA20 and hNAA25 are mediated between hNAA20 α1 helix and α26-α27 loop, and the hNAA25 α24-α25 loop, consisting of a mix of polar and non-polar interactions. A few hydrogen bonds are formed, centered around hNAA20-Asn16, between the sidechain nitrogen atoms of hNAA20-Asn16 and sidechain oxygen of hNAA25-Thr541, between the sidechain oxygen atom of hNAA20-Asn16 and sidechain nitrogen atom of hNAA25-Gln538, and between the sidechain nitrogen of hNAA20-Asn16 and the sidechain of hNAA25-Asp657 (**Fig. 3D**). A salt bridge is found between the sidechain of hNAA20-Asp10 and the sidechain of hNAA25-Arg602. In addition, there are hydrophobic interactions between residues Phe15, Leu12, Phe13, Leu19, Pro22 from hNAA20 and residues Ile537, Asp540, His536, Ala506, Phe662, Ser504, Ile542, Leu603 from hNAA25 (**Fig. 3D**).

Different sides of hNAA20 feature several potentially hNAA25-stabilizing polar interactions. Hydrogen bonds are formed between the Asp261 side chain of the hNAA25 α14-α15 loop and the hNAA20-Thr2 backbone nitrogen atom (**Fig. 3E**), between the Glu227 backbone carbonyl group of hNAA25 α12-α13 loop and the hNAA20-Arg102 sidechain (**Fig. 3F**), and between the Lys535 sidechain from NAA25 α26-α27 loop and the hNAA20-Ser141 backbone carbonyl group (**Fig. 3G**).

### hNatB makes specific interactions with the first 4 N-terminal residues of αSyn

In the cryo-EM map, density for the CoA-αSyn conjugate bisubstrate inhibitor is well resolved, allowing us to confidently model the CoA portion and the first five N-terminal residues (of ten residues present) of the αSyn portion (**Fig. 4A and 4B, Movie 2**). Similar to other NATs, CoA enters the catalytic active site through a groove formed by α3 and α4 of the catalytic subunit, while the peptide substrate enters the active site on the opposite side of the catalytic subunit flanked by the α1-2 and β6-β7 loops (**Fig. 4A)**.

**Fig. 4.**
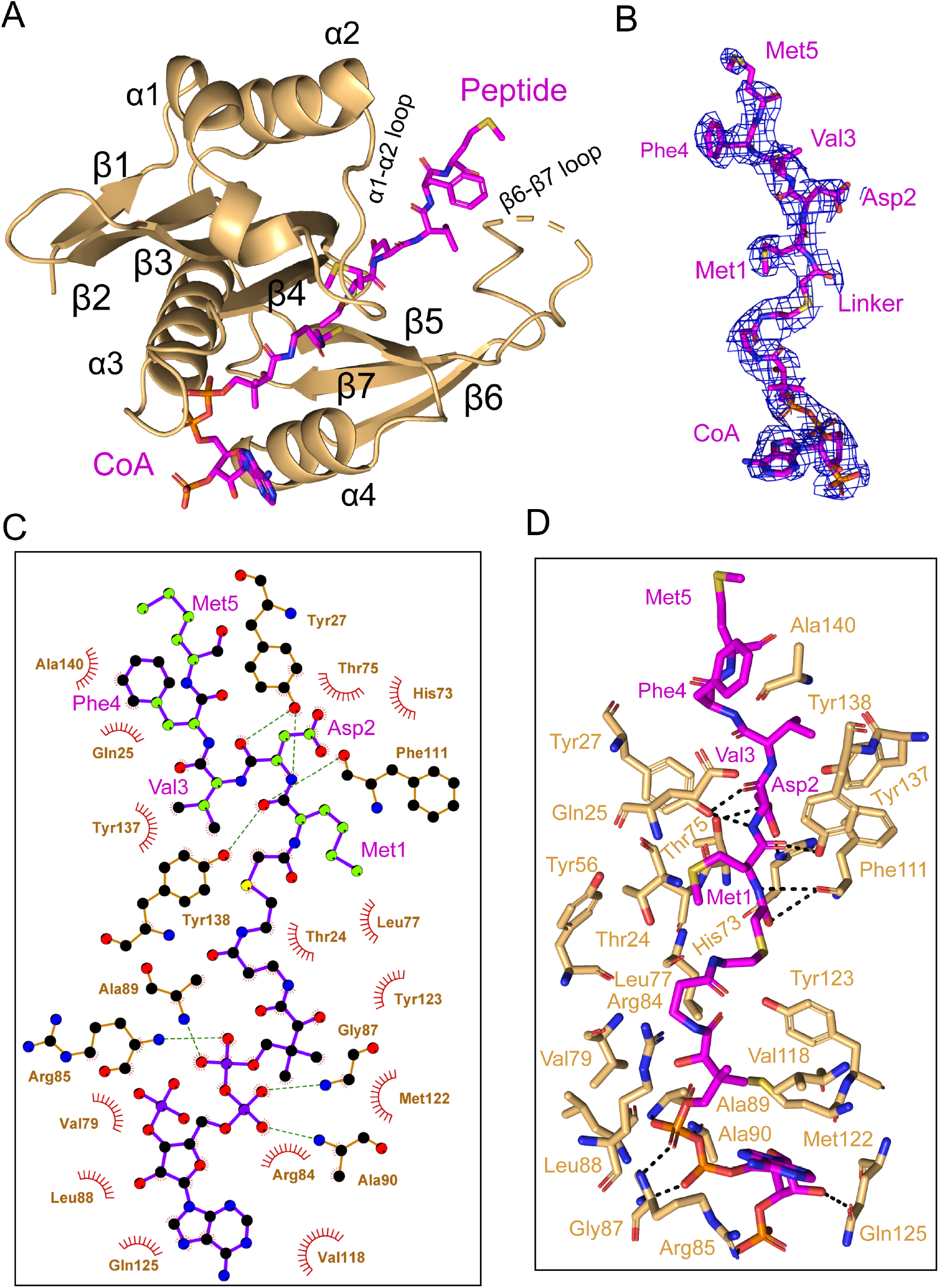
hNAA20 makes key CoA- and substrate peptide-interactions. **A**. The structure of hNAA20 bound to the CoA-αSyn conjugate bound is shown in cartoon, with corresponding secondary structures labeled. **B**. The fit of the CoA-αSyn conjugate ligand in the EM density map. The contour level is 4.0 sigma. **C**. Interaction between CoA-αSyn conjugate and hNAA20 residues is generated with LIGPLOT[80]. Hydrogen bonds are indicated by dashed green lines, and van der Waals interactions are indicated with red semicirlcles. **D**. Highlighted polar and hydrophobic interactions between CoA-αSyn conjugate and the hNAA20 are depicted in 3D view.

hNAA20 contains a conserved acetyl-CoA binding motif among NATs: R_84_R_85_XG_87_XA_89_ **(Fig. S5)**. Here, we observe that the positively charged hNAA20-Arg 85 interacts with the negatively charged 3’-phosphorylated ADP portion of CoA to form a salt bridge while Arg84 makes Van der Waals interactions (**Fig. 4C and 4D)**. A hydrogen bonding network is formed mainly between the 5’-diphosphate group and backbone atoms of a few residues including Val79, Gly87, Ala89, and Ala90 (**Fig. 4C and 4D)**, and mediated by the side chains of Arg85 and Gln125. The CoA molecule anchors to the binding pocket through a series of van der Waals contacts formed by residues Ser67, Val79, Leu77, Leu88, Val118, Met 122, and Tyr123 (**Fig. 4D)**.

Four N-terminal residues of αSyn participate in hNAA20 interactions. Anchoring of the αSyn peptide is mediated by protein hydrogen bonds with the backbone atoms of Met1 and Asp2 of αSyn. Hydrogen bonds are formed between the backbone N-H group of αSyn-Met1 and the backbone carbonyl group of hNAA20-Phe111, as well as the backbone carbonyl group of αSyn-Met1 with the sidechain of hNAA20-Tyr138. The backbone N-H and carbonyl of αSyn-Asp2 also form hydrogen bonds to the sidechain of Tyr27, and between the backbone carbonyl group of Asp2 and sidechain of hNAA20-Tyr27 (**Fig. 4D)**. Remarkably, hNAA20 contacts each of the first four N-terminal residue side chains of αSyn via van der Waals interactions. The only side chain that forms a hydrogen bond with hNAA20 is αSyn-Asp2, which hydrogen bonds with a hNAA20-His73 ring nitrogen and the hNAA20-Thr75 side chain (**Fig. 4D)**. The more extensive van der Waals interactions include the following: αSyn-Met1 interacts with hNAA20 residues Glu25, Phe27, Tyr56 and Ala76; αSyn-Asp2 interacts with hNAA20 residues Try27, Thr75, His73, Phe111, and Tyr138; αSyn-Val3 interacts with hNAA20 residues Tyr137 and Tyr138; and αSyn-Phe4 interacts with hNAA20 residues Glu25 and Als140. αSyn-Met5 does not appear to make specific interactions (**Fig. 4D)**. Consistent with the importance of the residues that mediate αSyn binding, most of the residues are highly conserved from yeast to humans **(Fig. S5)**

### Mutational analysis identifies key residues for hNatB catalysis and cognate substrate binding

To determine the functional importance of hNAA20 residues that appear to make important peptide or CoA substrate contacts in our model, we used an *in vitro* acetyltransferase assay to kinetically characterize WT and mutant hNatB proteins. Each mutant was purified to homogeneity and displayed identical gel filtration chromatography elution profiles (data not shown), indicating that they were all properly folded. We prepared alanine mutants of several residues involved in the CoA binding including Arg84, Arg85, Gly87 and Tyr123. Among them, R84A, R85A and G87A did not show significant defects in overall protein catalytic function (**Table 1**). However, a Y123A mutant nearly abolished protein activity, with a 95% loss of protein activity, affecting both *k_cat_* and K_m_ (**Table 1**). To further interrogate the properties of this residue, we prepared a Y123F mutant which features a similar aromatic bulky side chain but not the polar p-hydroxyl group. We observed that Y123F displayed a similar ∼88% loss of *k_cat_*, but had a negligible effect on the peptide K_m_ (**Table 1**). These data suggested that the Tyr123 hydroxyl group is critical for catalysis but not required for substrate binding, while the aromatic ring of Tyr123 plays a role in peptide substrate binding. Given that the hydroxyl group of Tyr123 is about 3.5 Å from the sulfur atom of the CoA-αSyn conjugate and 6.3 Å away from the αSyn N-terminus, it is in a position to play a role as a general base or acid for catalysis, potentially through an intervening water molecule **(Fig. 5A)**. This is analogous to the proposed general base role of Tyr31 as a general base for hNAA50 catalysis [31] (See discussion).

**Fig. 5.**
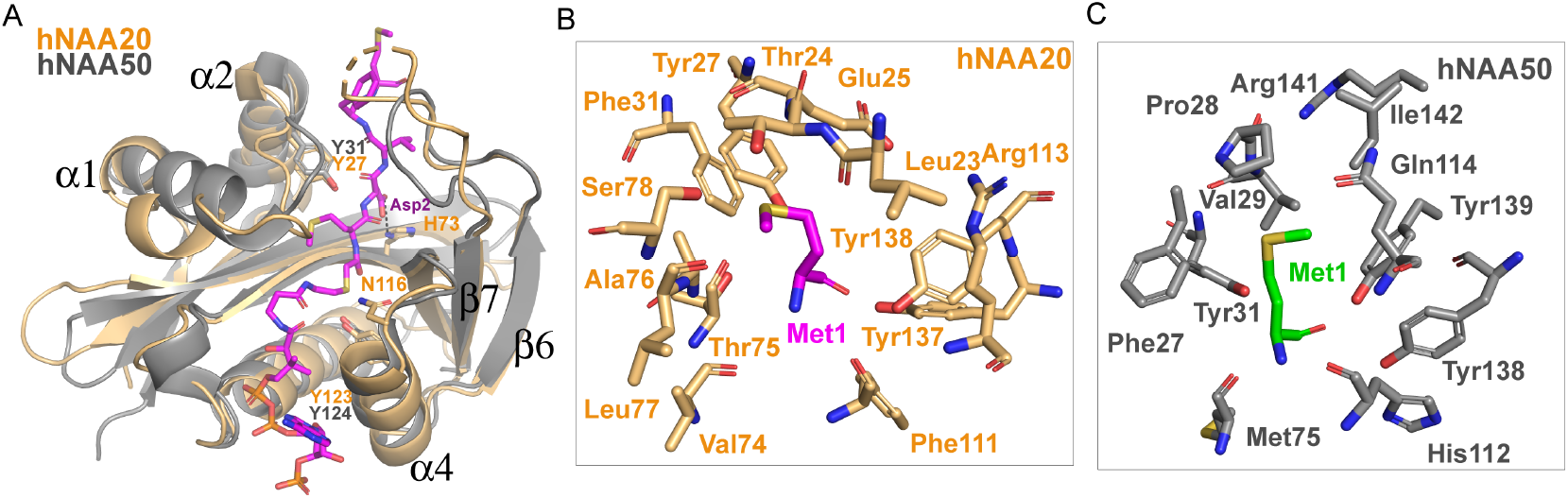
Structural comparison between hNAA20 and hNAA50. **A**. Bi-substrate inhibitor-bound hNAA20 (light orange) is shown superimposed with hNAA50 (grey, PDB: 3TFY). H73, N116, and Y123 (sticks), mediate important functional roles in hNatB catalysis. **B**. Residues forming the Met1 binding pocket of hNAA20 are depicted. **C**. Residues forming the Met1 binding pocket of hNAA50 are depicted.

We also prepared alanine substitutions for residues that appeared to play important roles in αSyn binding: Glu25, Tyr27, His73, Tyr137, Tyr138. We were surprised to find that mutations of hNatB residues that mediated backbone hydrogen bond interactions, Y27A and Y138A, had relatively modest effects on αSyn peptide NTA with Y27A showing ∼2-fold higher K_m_ and Y138A showing ∼2-fold reduced K_cat_, together suggesting that side chain contacts might dominate the binding energy (**Table 1**). Consistent with this, and our structural observations, we found that H73A produced a ∼90% reduction in activity (**Table 1**). This correlates with the importance of the His73 hydrogen bond and van der Waals contacts with αSyn-Asp2. Of note, other cognate hNatB side chain residues at position 2, Glu, Gln, and Asn would also be well positioned to form hydrogen bonds with His73. Together, hNAA20-His73 appears to play a critical role in cognate substrate recognition by hNatB.

hNatB-Asn116 is a highly conserved NatB residue **(Fig. S5)** that caps the hNAA20 a4 helix, which also harbors the putative catalytic residue, Tyr123, and is also in position to make a water-mediated hydrogen bond with to CoA pantetheine nitrogen **(Fig. 5A)**. This observation suggests that Asn116 could play an important functional role. To test this, we prepared and evaluated an N116A mutant and, consistent with our hypothesis, Ans116, we found that this mutant leads to ∼ 90% loss in activity (**Table 1**).

Taken together, our structural and mutational analysis of hNatB highlight the functional importance of hNatB-NAA20 residues His73, Asn116, and Tyr123 in hNatB-mediated N-terminal acetylation. While Tyr123 appears to play a critical catalytic role, potentially as a general base and/or acid for catalysis; His73 appears to play an important role in the recognition of substrate residue 2 and Asn116 likely plays a structural role **(Fig. 5A)**. Each of these residues could employ a bridging water molecule to mediate their functional roles, although these putative water molecules are not visible at the current resolution of our structure.

## Discussion

Since the identification of hNatB more than a decade ago, many studies have shown that it N-terminally acetylates important proteins such as actin, tropomyosin, CDK2, and α-Synuclein, and its function has connections to diseases such as hepatocellular carcinoma and Parkinson disease [17, 42, 43, 45, 54, 55]. Despite its clear biological importance, hNatB-mediated NTA by hNatB remained poorly understood. Here, we developed a CoA-αSyn conjugate hNatB inhibitor, determined the cryo-EM structure of CoA-αSyn inhibitor-bound hNatB, and carried out associated structure-guided mutagenesis and activity assays. This has led to the identification of functionally important differences with human NatA and *C. albicans* NatB. These studies have also provided evidence for important hNatB-specific elements responsible for αSyn recognition and N-terminal acetylation, providing direct implications for NatB recognition of other canonical substrate proteins.

Consistent with previous studies, we have demonstrated that hNatB acetylates a cognate “MD” N-terminus, and is unable to N-terminally acetylate non-cognate N-termini that are substrates for other NATs such as NatA, NatC, NatE. We demonstrated for the first time that hNatB can acetylate an α-Syn peptide *in vitro*, directly linking hNatB to Parkinson disease. We have also demonstrated that αSyn peptides lacking the Met1 or both the Met1 and Asp2, do not serve as hNatB substrates, confirming the strict substrate specificity of hNatB. This is consistent with our structural model showing significant interactions between hNatB-NAA20 and both the first and second N-terminal residues of an αSyn peptide, with important but less extensive interactions with the third and fourth residues. This hierarchy of interactions likely explains how NatB enzymes can accommodate cognate substrates that diverge at positions three and four.

Here, we have presented hNatB-αSyn interactions that can be used to rationalize the substrate specificity of hNatB: N-terminal sequences containing “MD-”, “ME-”, “MN-”, and “MQ-”. αSyn-Met1 sits in a hydrophobic pocket that comfortably accommodates a methionine residue, whereas shorter side chains or longer polar or charged side chains would fit poorly (**Fig.5B-C**). The nature of this binding pocket is similar to the described hNAA50 recognition of Met1 [31] (**Fig. 5B-C**). Although both of hNAA50 and hNAA20 can N-terminally acetylate peptides with Met at the first position, no overlapping activity has been observed. This can be rationalized based on the chemical properties of the second residue in the cognate peptide. We find that the αSyn-Asp2 sidechain forms hydrogen bonds with the hNAA20 side chains His73 and Thr75. These polar residues in the peptide binding site of hNAA20 would likely serve as poor acceptors of the largely hydrophobic residues targeted by hNAA50. The hNatB substrate client profile featuring D-, E-, N- or Q-residues in position two is consistent with the mechanisms of substrate recognition observed in the binding pocket for αSyn-Asp2. The aliphatic regions of each of these side chains (D, E, N, Q) would all benefit from the extensive hNatB van der Waals interactions surrounding the aliphatic region for αSyn-Asp2 (Tyr27, His73, Thr75, Phe111, and Tyr138). This second residue would also form a hydrogen bond interaction with His73 and Thr75, which may accommodate the carboxyl side chains of both the shorter D- and N- and longer E- and Q-side chains. Notably, His73 and Thr75 are strictly conserved from yeast to man **(Fig. 5 and Fig. S3)**. In contrast, shorter polar or nonpolar side chains would less efficiently fill the pocket for residue two, while larger polar or charged side chains would likely result in steric clashes. In agreement with this, we have demonstrated that H73A mutation has severe impact on hNatB catalysis (**Table 1**).

The hNatB/CoA-αSyn structure has implications for the mode of hNatB catalysis. While previous studies have suggested that the general base for hNAA50 catalysis is hNAA50-Tyr31, mutation of the corresponding hNaa20 residue, Tyr27, had minimal effects on hNatB kinetic parameters. Strikingly, our mutational analysis has identified the functional importance of hNatB-NAA20 residues His73, Asn116 and Tyr123, although Tyr123 is the only residue that is in position to play a catalytic role (**Fig. 5A**). Specifically, Tyr123 is in position to play a catalytic role, potentially as a general base and/or acid, through a bridging water molecule (although a water molecule is not visible at the current resolution). Interestingly, hNAA50 contains a tyrosine residue at the same position (hNAA50-Tyr124), although the mechanistic significance of this tyrosine residue has not yet been described [31]. It would be of interest to determine if hNAA50-Tyr124 also plays an important catalytic role (possibly in combination with hNAA50-Tyr31), similar to the corresponding hNAA20-Tyr123 of hNatB.

The biological importance of hNatB and its connection to various disease processes highlights it as an important target for probe and inhibitor development. Indeed, a recent study highlights hNatB as a therapeutic target for αSyn toxicity [56]. Our development of a CoA-αSyn conjugate bisubstrate with an IC_50_ of ∼ 1.6 μM represents a step in this direction, although the structural information provided here could further aid to the rational development of more drug-like hNatB inhibitors with possible therapeutic applications.

## Materials and Methods

### Protein Expression and Purification

hNAA20 with a C-terminal truncation (1-163 out of 178 residues) and full-length hNAA25 were cloned into two separate insect cell expression vectors pFASTBac HTA. hNAA20 was untagged, while hNAA25 contained a Tobacco-etch virus (TEV)-cleavable N-terminal 6xHis-tag. Human NatB complex (hNAA20^1-163^/hNAA25^FL^) was obtained by coexpressing these two plasmids in Sf9 (*S. frugiperda*) cells (ThermoFisher, cat# 12659017), and purified as described previously[35]. Sf9 cells were grown to a density of 1×10^6^ cells/ml and infected using the amplified hNAA20^1-163^/hNAA25^FL^ baculovirus to an MOI (multiplicity of infection) of 1-2. The cells were grown at 27 °C and harvested for 48 hours post-infection by centrifugation. Cell pellets were resuspended in lysis buffer (25 mM Tris, pH 8.0, 300 mM NaCl, 10 mM Imidazole, 10 mM β-ME, 0.1 mg/mL PMSF, DNase, and complete, EDTA-free protease inhibitor tablet) and lysed by sonication. After centrifugation, the supernatant was isolated and passed over Ni-NTA resin (Thermo Scientific), which was subsequently washed with 10 column volumes of lysis buffer. Protein was eluted with a buffer with 25 mM Tris, pH 8.0, 300 mM imidazole, 200 mM NaCl, 10 mM β-ME, which was dialyzed into buffer with 25 mM HEPES pH 7.5 50 mM NaCl 10 mM β-ME. Ion-exchange was carried out with an SP ion-exchange column (GE Healthcare) in dialysis buffer with a salt gradient (50-750 mM NaCl). Peak fractions were concentrated to ∼ 0.5 mL with a 50-kDa concentrator (Amicon Ultra, Millipore), and loaded onto an S200 gelfiltration column (GE Healthcare) in a buffer with 25 mM HEPES, pH 7.5, 200 mM NaCl, and 1 mM TCEP. Proteins were aliquoted, snap-frozen in liquid nitrogen, and stored at −80 °C for further use. Protein harboring mutations were generated with the QuickChange protocol (Stratagene) and obtained following the same expression and purification protocol as described for the wild-type protein. Primers synthesized for the generation mutant constructs are listed in the Supplementary file.

### Acetyltransferase Activity Assays

All acetyltransferase assays were carried out at room temperature in a reaction buffer containing 75 mM HEPES, pH 7.5, 120 mM NaCl, 1 mM DTT as described [33, 34]. The “MDVF” peptide substrate was based on the first 7 amino acid of α-Synuclein (“MDVF” peptide: NH_2_-MDVFMKGRWGRPVGRRRRP-COOH; “SASE” peptide: NH_2_-SASEAGVRWGRPVGRRRRP-COOH; “MLGP” peptide: NH_2_-MLGPEGGRWGRPVGRRRRP-COOH; “SGRG”/H4 peptide: NH_2_-SGRGKGGKG LGKGGAKRHR-COOH; “MLRF” peptide: NH_2_-ML RFVTKRWGRPVGRRRRP-COOH; “DVFM” peptide: NH_2_-DVFMKGLRWGRPVGRRRRP-COOH; “VFMK” peptide: NH_2_-VFMKGLSRWGRPVGRRRRP-COOH; GenScript). Reactions were performed in triplicate. To determine steady-state catalytic parameters of hNatB with respect to acetyl-CoA, 100 nM hNatB was mixed with a saturating concentration of “MDVF” peptide substrate (500 μM) and varying concentrations (1.95 μM to 1 mM) of acetyl-CoA (^14^C-labeled, 4 mCi mmol^−1^; PerkinElmer Life Sciences) for 10-minute reactions. To determine steady-state catalytic parameters of hNatB with respect to peptide substrate, 100 nM hNatB was mixed with saturating concentrations of acetyl-CoA (300 μM, ^14^C-labeled) and varying concentrations of “MVDF” peptide (1.95 μM to 1 mM) for 10 minutes. Reactions were quenched by adding the solution to P81 paper discs (Whatman). Unreacted acetyl-CoA was removed by washing the paper discs in buffer with 10 mM HEPES, pH 7.5, at least three times, each 5 minutes. The paper discs were then dried with acetone and transferred to 4 mL scintillation fluid for signal measurement (Packard Tri-Carb 1500 liquid scintillation analyzer). Data was fitted to a Michaelis–Menten equation in GraphPad Prism to calculate kinetic parameters. Kinetic parameters on mutants with respect to peptide were carried out in the same condition as for wild type, with 300 μM ^14^C labeled acetyl-CoA and varied peptide concentration (1.95 μM to 1 mM). All radioactive count values were converted to molar units with a standard curve created with known concentrations of radioactive acetyl-CoA added to scintillation fluid. GraphPad Prism (version 5.01), was used for all data fitting to the Michaelis–Menten equation. For IC_50_ determination of the CoA-αSyn conjugate, 100 nM hNatB was mixed with 500 μM “MVDF” peptide and 300 μM ^14^C labeled acetyl-CoA, and inhibitor concentrations were varied (0.23 μM to 13.44 μM). Data were fit to a sigmoidal dose-response curve with GraphPad Prism (version 5.01). Errors represent s.d. (n = 3).

### Cryo-EM data collection

For initial sample screening, 0.6 mg/ml fresh hNatB sample with three-molar excess bisubstrate was used. hNatB particles on these grids exhibited a severe preferred orientation, which generated an incorrect 3D initial model (data not shown). To solve this issue, 1 μL of 0.05% NP-40 was mixed with 20 μL of hNatB (4 mg/mL). 3 μL of this sample was applied to Quantinfoil R1.2/1.3 holey carbon support grids, blotted and plunged into liquid ethane, using an FEI Vitrobot Mark IV. An FEI TF20 was used for screening the grids and data collection was performed with a Titan Krios equipped with a K3 Summit direct detector (Gatan).

### Cryo-EM data processing

Original image stacks were summed and corrected for drift and beam-induced motion at the micrograph level using MotionCor2 [69]. Defocus estimation and the resolution range of each micrograph were performed with Gctf [70]. About 3000 particles were manually picked to generate several rough 2D classaverages. Representative 2D classes were used to automatically pick ∼1,927,673 particles from 5281 micrographs in Relion 3.0 [71]. All particles were extracted and binned to accelerate the 2D and 3D classification. After bad particles were removed by 2D and 3D classification, 982, 420 particles were used for auto-refinement and per particle CTF refinement. The final map was refined to an overall resolution of 3.46 Å, with local resolution estimated by Resmap [72].

### Cryo-EM model building and refinement

The hNatB atomic model was manually built de novo using the program COOT [73] according to the cryo-EM map, with the guidance of predicted secondary structure and bulky residues such as Phe, Tyr, Trp and Arg. The first two alpha helices of hNAA25 were built as poly-alanine, due to the lack of tracible density in the 3D map. The complete model was then refined by real-space refinement in PHENIX [74]. All representations of cryo-EM density and structural models were performed with Chimera [75] and PyMol (https://pymol.org/2/). The sequence alignments with secondary structure display were created by ESPript 3.0 [76]. hNAA25 TPR predictions were performed using the TPRpred server [77, 78] (https://toolkit.tuebingen.mpg.de/#/tools/tprpred). The surface area calculation was performed using PDBePISA [79] (Proteins, Interfaces, Structures and Assemblies) (http://www.ebi.ac.uk/pdbe/pisa/).

## ACKNOWLEDGEMENTS

This work was supported by NIH grant R35 GM118090 awarded to R.M. and R01 NS103873 awarded to E.J.P. B.P. thanks the University of Pennsylvania for support through a Dissertation Completion Fellowship. We acknowledge the support of the Perelman School of Medicine, University of Pennsylvania DNA Sequencing Core Facility and D. Schultz and E. Dean from the University of Pennsylvania High Throughput Screening Core Facility for providing the Sf9 cells expressing hNatB for this study. We thank Dr. Zuo Biao and Dr. Sudheer Molugu from the University of Pennsylvania Electron Microscopy Resource Lab for help with initial cryo-grid screening; and Dr. Darrah Johnson-McDaniel from the Beckman Center for Cryo-EM at the University of Pennsylvania for technical assistance on data collection. We also thank Dr. Xuepeng Wei for help in data analysis and discussion.

## Author Contribution

Conceptualization, S.D., B.P., L.G., E.J.P. and R.M.; Methodology, S.D., B.P., L.G., E.J.P.; Investigation, S.D., B.P., and L.G.; Formal Analysis, S.D; Writing – Original Draft, S.D.; Visualization, S.D.; Writing – Review and Editing, S.D., B.P., L.G., E.J.P. and R.M.; Funding Acquisition, R.M. and E.J.P.; Resources, R.M.; Supervision, R.M. and E.J.P..

## Supplementary information

**Supplementary Table 1.**
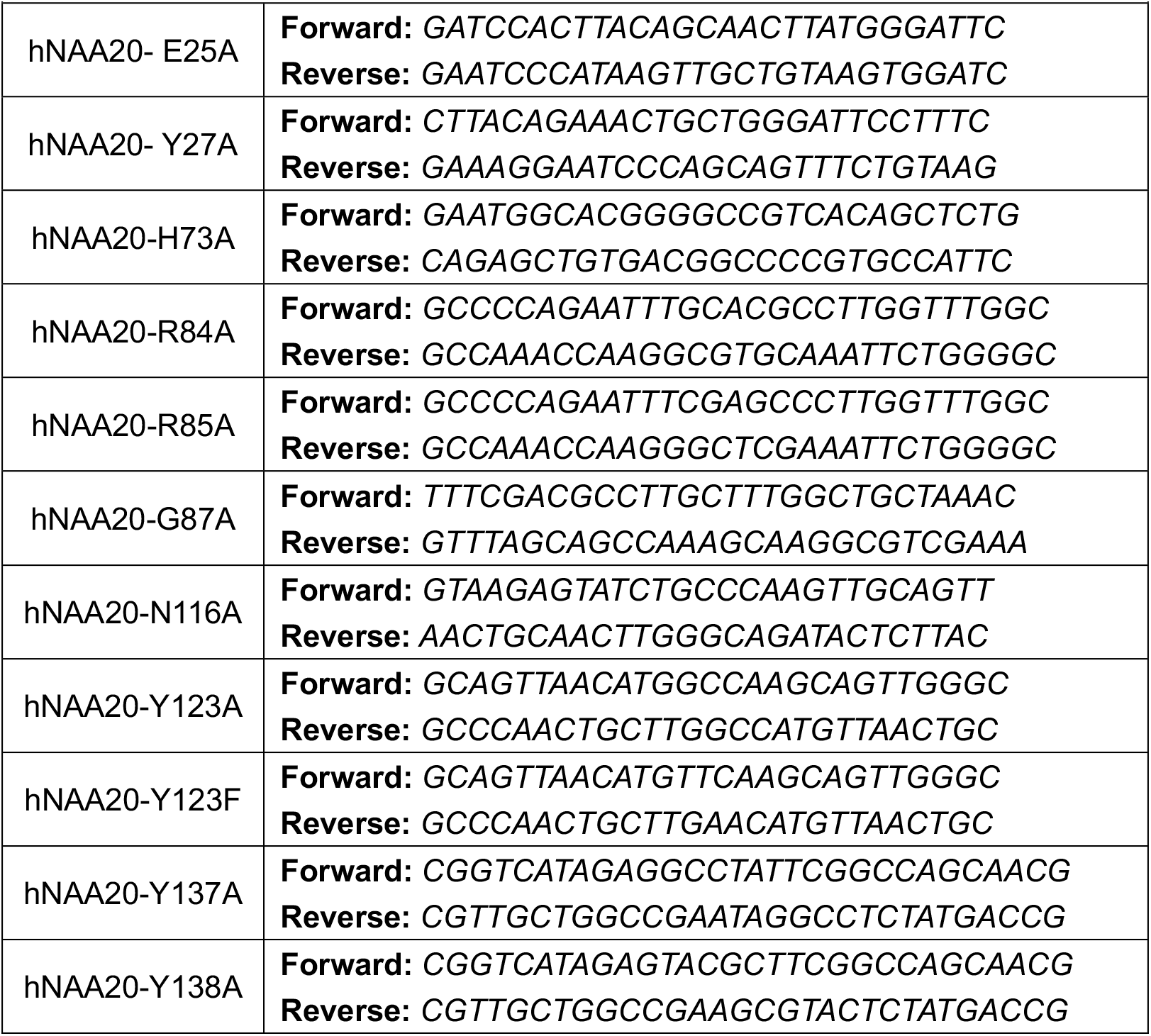
Primers sequence for preparing mutations.

**Fig. S1.**
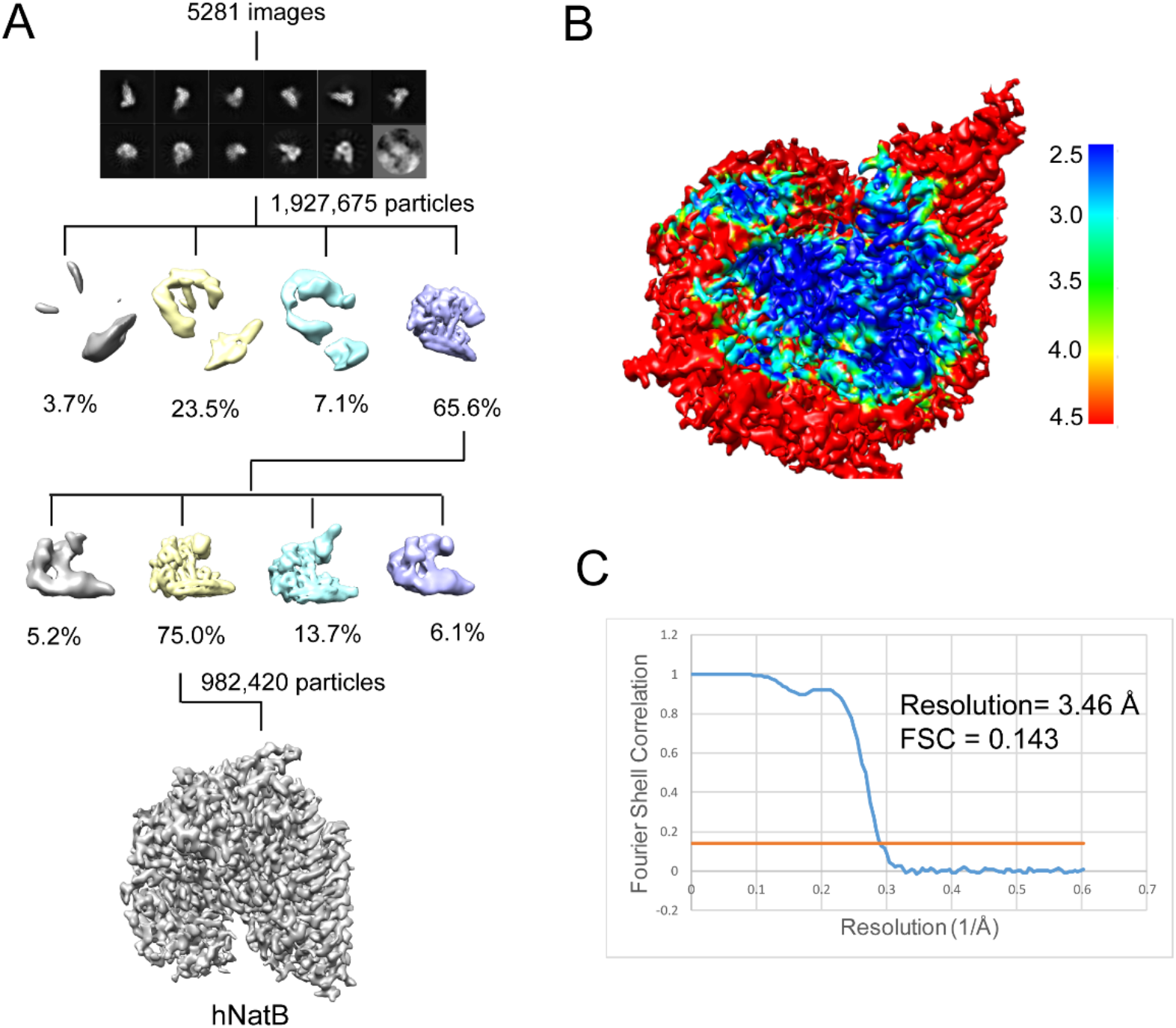
Cryo-EM workflow and resolution of hNatB map. **A**. 2D and 3D classification scheme for hNatB EM map determination. **B**. Local resolution map of hNatB. **C**. FSC curve of hNatB EM map.

**Fig. S2.**
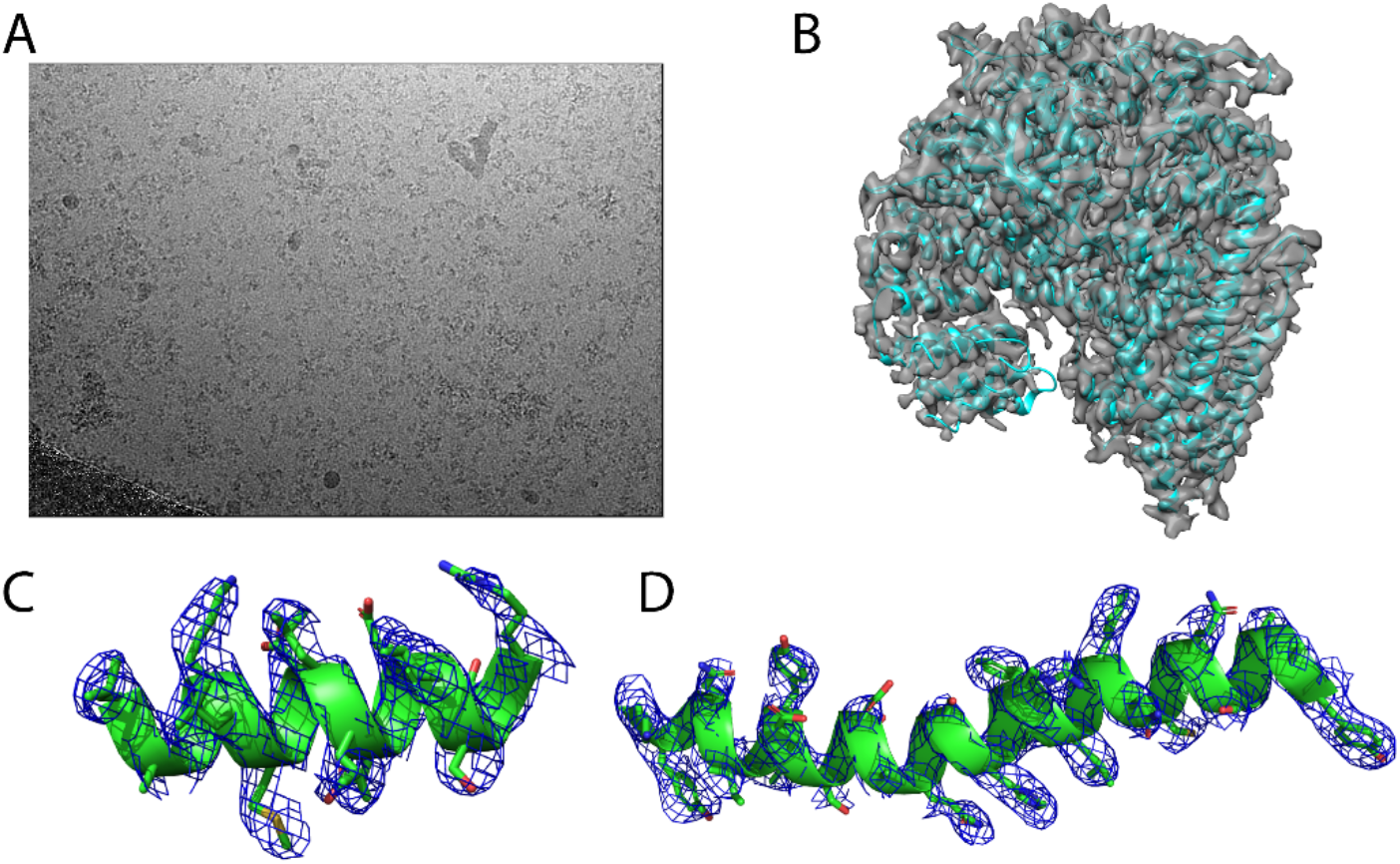
Representative micrograph of hNatB cryo-images and hNatB model fit-in map. **A**. A representative micrograph of a hNatB frozen grids hole **B**. Atomic model of hNatB fitted into the Cryo-EM map. **C**. The fit of a helical segment from hNAA20 in the EM density. The contour level is 5 sigma. **D**. The fit of a helical segment from hNAA25 in the EM density. The contour level is 5 sigma.

**Fig. S3.**
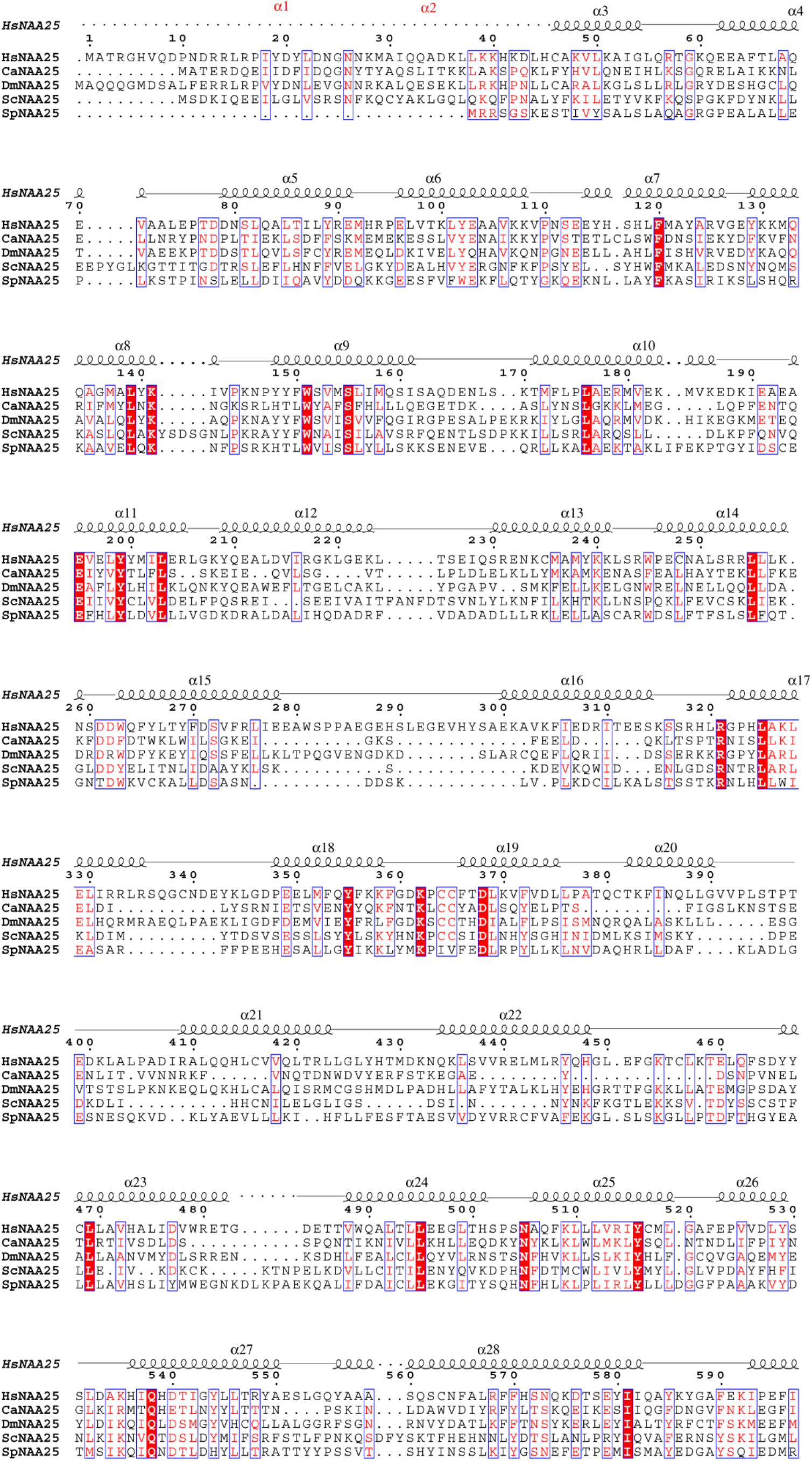
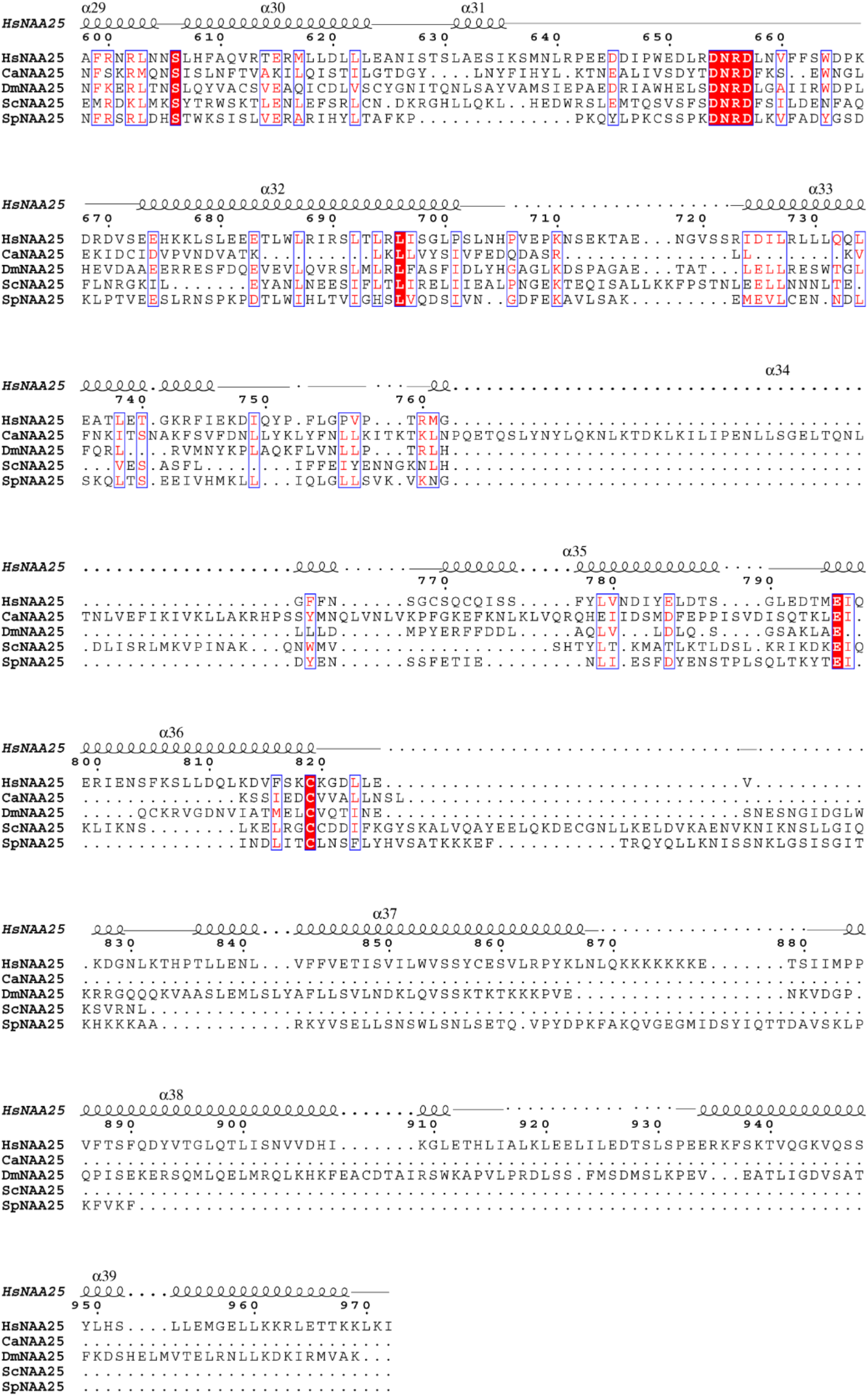
Sequence alignment of NAA25 homologues. Species include *H. sapiens* (Hs), *C. albicans* (Ca), *D. melanogaster* (Dm), *S. cerevisiae* (Sc). and *S. pombe* (Sp).

**Fig. S4.**
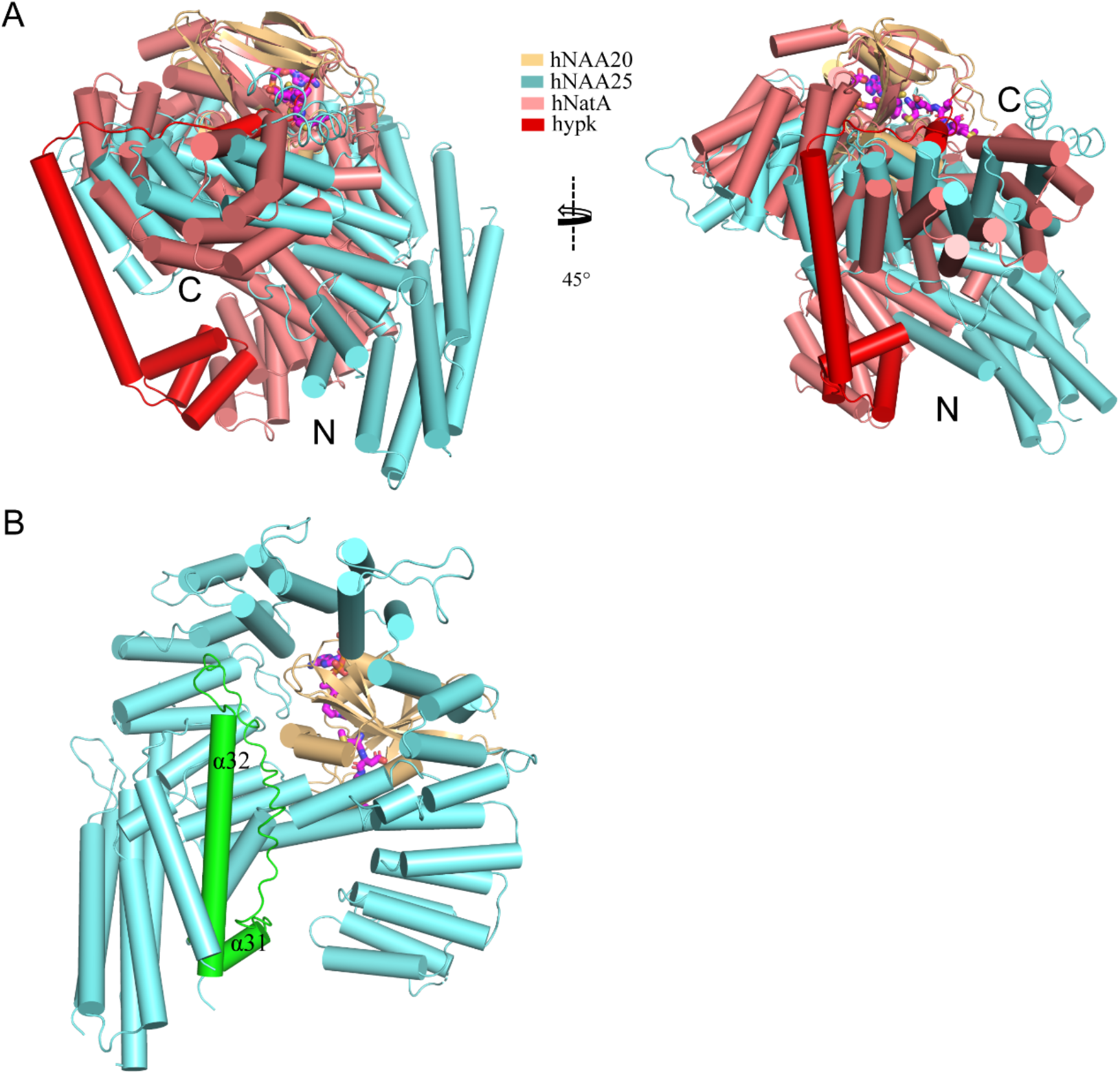
Structural comparison of hNatB and hNatA-HYPK. **A**. hNatB overlaid with hNatA-HYPK. hNatA and HYPK are colored as light salmon and red, respectively. N- and C-terminiof the auxiliary subunits are indicated. **B**. The extended loop connecting α31 to α32 is highlighted in green in hNatB. This loop is not present in hNatA (not shown).

**Fig. S5.**
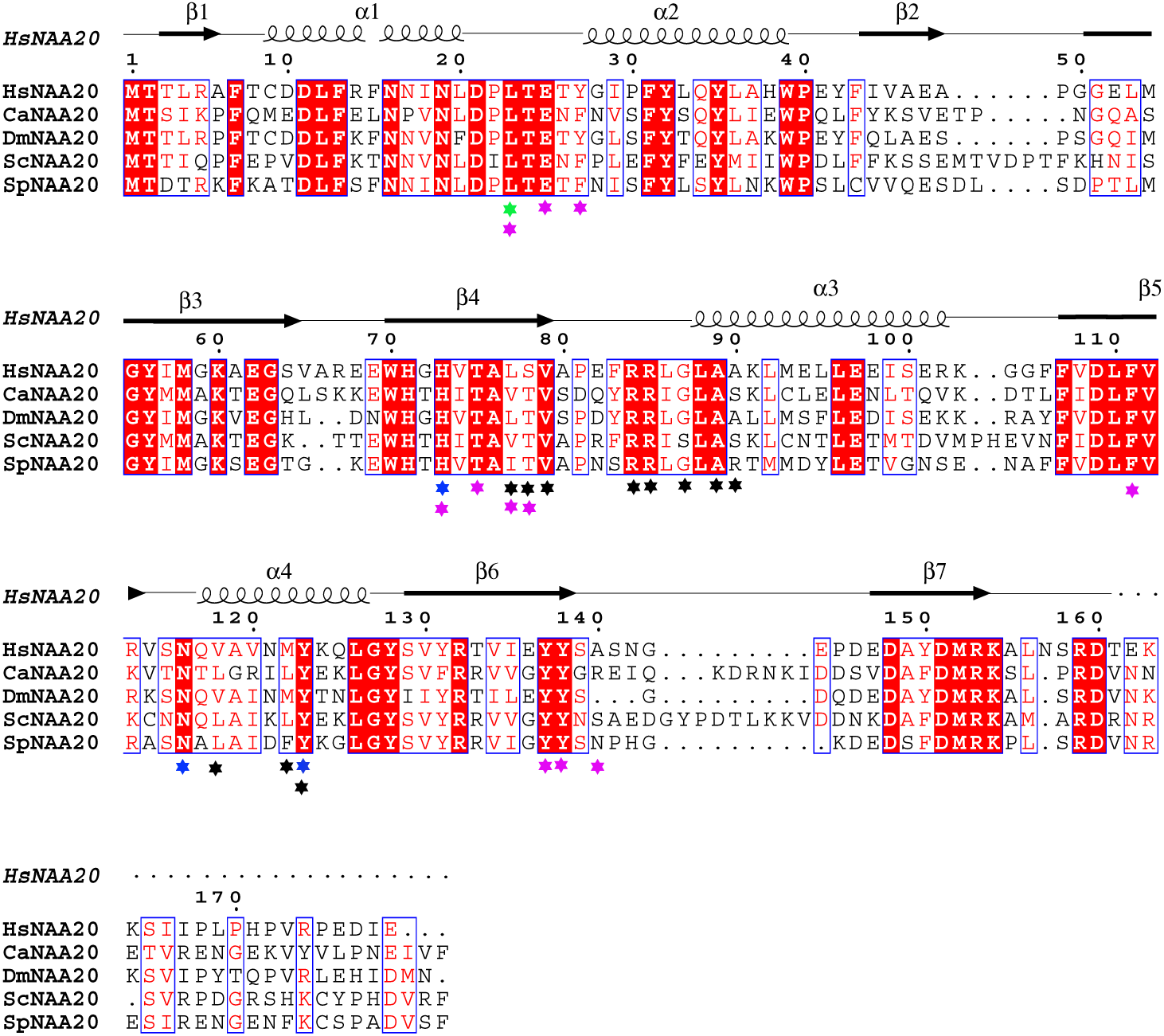
Sequence alignment of NAA20 homologues. Species depicted include *H. sapiens* (Hs), *C. albicans* (Ca), *D. melanogaster* (Dm), *S. cerevisiae* (Sc). and *S. pombe* (Sp). Blue, black and magenta labeled indicate mutation sensitive residues, CoA-binding residues, and peptide-binding residues, respectively.

## Movie Legends

**Movie 1. Overall view of the NatB complex**.

hNaa20 (light orange) and hNaa25 (cyan) are shown in cartoon. The CoA-αSyn conjugate inhibitor is shown in sticks and colored as magenta.

**Movie 2. Overall view of α-synuclein N-terminal interactions by NAA20**.

Amino acid sidechains that mediate hydrogen bond and van der Waals interactions with a-synuclein are highlighted on a cartoon model of NAA20.

